# bettercallsal: better calling of *Salmonella* serotypes from enrichment cultures using shotgun metagenomic profiling and its application in an outbreak setting

**DOI:** 10.1101/2023.04.06.535929

**Authors:** Kranti Konganti, Elizabeth Reed, Mark Mammel, Tunc Kayikcioglu, Rachel Binet, Karen Jarvis, Christina M. Ferreira, Rebecca L. Bell, Jie Zheng, Amanda M. Windsor, Andrea Ottesen, Christopher Grim, Padmini Ramachandran

## Abstract

Precise and rapid identification of *Salmonella* serotypes from suspect food matrices is critical for successful source attribution of illness outbreaks (Scallan et al., 2011). Currently, close to 3% of U.S. foodborne *Salmonella* outbreaks have been attributed to multiple S*almonella* serotypes (2.85%, 2000 - 2020) (CDC, 2022). Recent foodborne outbreaks that have been attributed to multiple *Salmonella* serotypes force us to question whether these are rare events or if previous methods simply did not have the throughput to provide an accurate picture of the complex ecology that is connected to outbreak etiologies. (Hassan et al., 2019; FDA, 2021; Whitney et al., 2021).

An *in-silico* benchmark dataset, comprising 29 unique *Salmonella*, 46 non-*Salmonella* bacterial and 10 viral genomes, was generated with varying read depths. For outbreak samples, analysis was performed on previously sequenced pre-enrichments and selective enrichments of papayas and peaches (fruits and leaves) that led to the identification of multiple serovars. Data analyses was performed using a custom-built *k-mer* tool, SeqSero2, Kallisto and bettercallsal.

The *in-silico* dataset analyzed with bettercallsal had accuracy, recall and specificity of 95%, 96% and 98 % respectively. In the papaya outbreak samples, bettercallsal identified multiple serovar presence in concordance with Bioplex assay results and the genome hits assigned to the samples are *Salmonella* isolates from the papaya outbreak as evident by NCBI SNP cluster information. In peach outbreak samples, bettercallsal identified both the serovars (Alachua and Gaminara) in concordance with *k*-mer analysis and the Luminex xMap assay. bettercallsal outperformed *k-mer*, Kallisto and Seqsero2 in identifying multiple serovars from enrichment cultures using shotgun metagenomics sequencing.

Most *Salmonella* subtyping work has relied upon WGS methods which focuses on the high-resolution analysis of single genomes, or multiple single genomes picked from colonies on agar. Here we introduce laboratory and bioinformatics innovations for a metagenomic outbreak response workflow that accurately identifies multiple *Salmonella* serovars at the same time in a much higher throughput approach. bettercallsal is one of the first analysis tools that can potentially identify multiple *Salmonella* spp. serotypes from a metagenomic or quasi-metagenomic datasets with accuracy and can provide early insights into the etiology of the sample.

## Introduction

*Salmonella* is one among the leading bacterial causes of foodborne outbreaks in the United States (Scallan et al., 2011). Understanding the relative public health impact of microbiological hazards across the food supply is critical for the food safety system. Current surveillance for *Salmonella* is limited to the detection of only the most abundant serotype(s) due to the use of culture-based approaches. Thus, some serotypes that are present in low abundance may remain undetected and, in some cases, even the etiological agent causing the outbreak remains undetected despite epidemiological connections to the food being tested. Next-generation sequencing (NGS) technologies have ushered in an era of precision analysis, transforming the way we detect, identify, and conduct source-tracking of foodborne pathogens. As we continue to discover more information about outbreak etiology, the potential of quasi-metagenomic methods for rapid detection is becoming abundantly clear (Ottesen et al., 2016; Ottesen et al., 2020).

The accurate subtyping and subsequent clustering of *Salmonella* strains associated with a foodborne outbreak event is essential for successful investigation and traceback to a specific food or environmental source. Most of the metagenomic profiling tools using either marker-based or *k-mer* based approaches for classification are sensitive down to the species rank but not to the strain level (McIntyre et al., 2017) and cannot accurately discern between the highly clonal Salmonella spp. serotypes. Assembly based approaches are also becoming widely used in metagenomics, especially when we want to obtain cluster information for an efficient traceback (Buytaers et al., 2021). The recent advent of DNA sketching based algorithms have enabled much more efficient and accurate processing of huge amounts of data with the potential of analyzing the sequencing data in “real-time” (Rowe, 2019). In the case of quasi-metagenomic datasets, we have in the past used CFSAN SNP pipeline successfully for clustering and traceback (Ottesen et al., 2020), but the pipeline performs best when there is appreciable coverage at all possible sites and when there is a single etiologic agent (Davis et al., 2015). Those criteria are difficult to achieve on every quasi-metagenomic dataset, and highly dependent on the food matrix and the pathogen of interest. Using approaches like *k-mer* or assembly-based methods or adapting tools that are applicable to whole genome sequencing on a multi-serovar outbreak have not resulted in detection of all the serotypes and the clustering information for a traceback. We have built bettercallsal to address the need to identify multiple *Salmonella* spp. serotypes from metagenomic or quasi-metagenomic datasets. We leverage the NCBI Pathogen Detection (PD) project and provide the link outs to the isolate genome hits via the NCBI Isolates Browser which in turn links out to the NCBI SNP Tree Viewer if that genome hit is a member of a clonally related cluster (Sayers et al., 2021).

This approach can have a true impact on understanding differences in the dynamics of individual *Salmonella* strains with different serotype and can have a significant impact on understanding the ecology of this pathogen with respect to food safety and public health measures. The bettercallsal workflow is licensed under MIT and is freely available for download and use from https://github.com/CFSAN-Biostatistics/bettercallsal.

## Materials and Methods

### Simulated datasets

Three sets of *in-silico* Illumina datasets were generated with InSilicoSeq (Gourle et al., 2019) using *Salmonella* and non-*Salmonella* microbial genome assemblies. The first dataset comprises 29 *Salmonella* genomes representing unique *Salmonella* serotypes of importance in foodborne illnesses along with 46 non-*Salmonella* bacterial and 10 viral and phage species (Supplementary table 1). This read set was generated at an equal coverage of 5X (sal-cov5x) for each of the 29 unique *Salmonella* serotypes and with a coverage of 0-12X for rest of the genomes (Supplementary table 1). This same composition of 85 genomes were used to generate the second read set with the only change being that the coverage of the 29 unique *Salmonella* genomes was between 1X to 5X (sal-cov1-5x) (Supplementary table 1). A third read set was generated with a log-normal abundance distribution for all the 85 genomes using the same random seed (`--seed` 27) at varying read depths ranging from 0.5 to 5 million read pairs using the MiSeq error model (Supplementary table 1). Additionally, to mimic some of the recently identified multi-serovar *Salmonella* serotypes isolated from a single sample linked to papaya outbreaks (Whitney et al., 2021), we generated 4 additional simulated Illumina paired-end datasets using InSilicoSeq (Gourle et al., 2019) with the MiSeq error model. Each dataset consisted of a mixture of 3 to 5 unique *Salmonella* spp. serotypes mixed in with the same 46 non-*Salmonella* bacterial species, and 10 viral and phage species (mix1 to mix4) (Table 1 and Supplementary table 1).

**Table 1:**
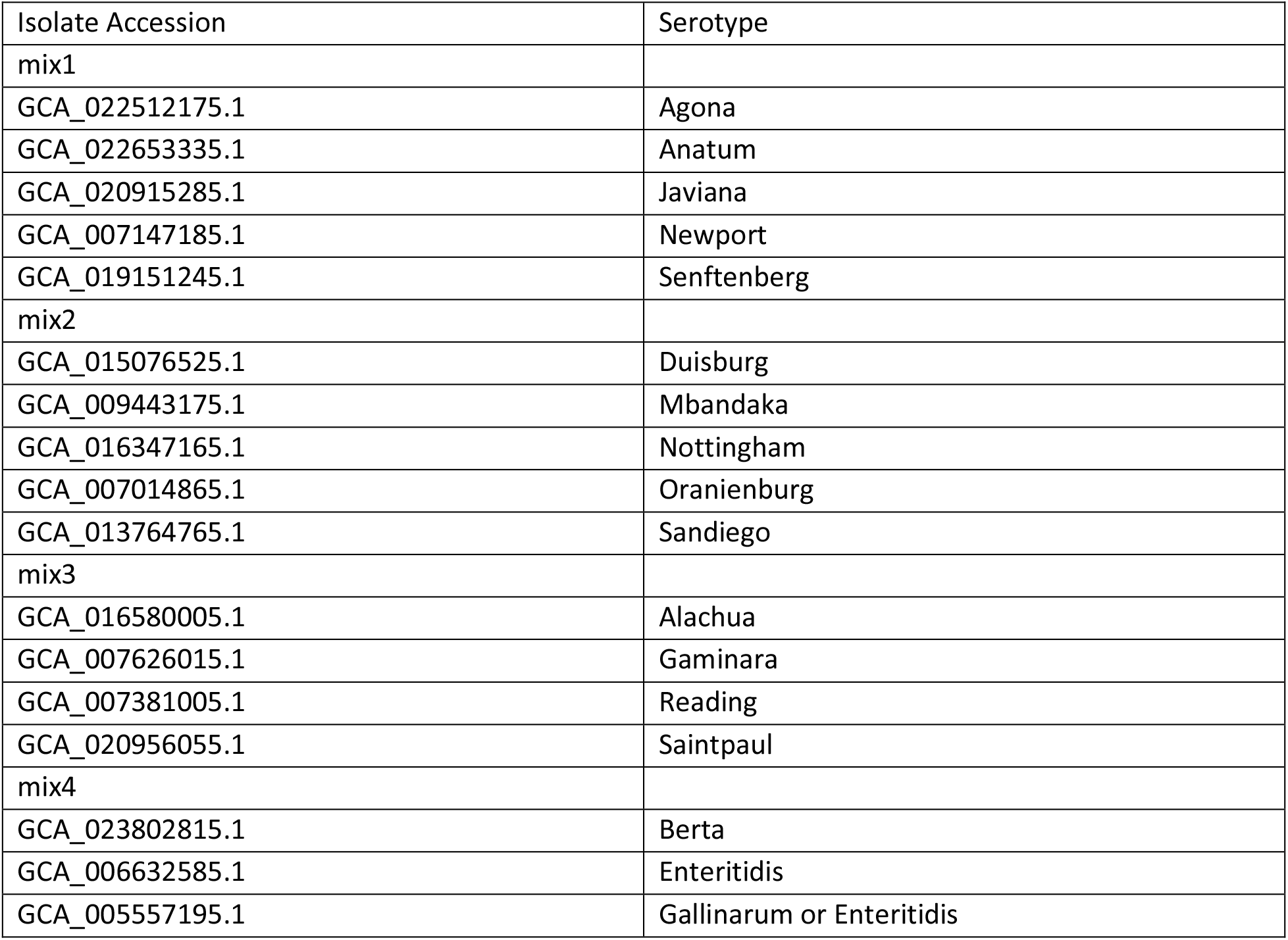
The *Salmonella* spp. serotypes in each of the mix1 to mix4 simulated datasets. InSilicoSeq was used with `- -seed 27` and MiSeq error model. The other 56 non-*Salmonella* genome accessions that are part of each of the mixes are described in Supplementary table 1.

| Isolate Accession | Serotype |
| --- | --- |
| mix1 |  |
| GCA_022512175.1 | Agona |
| GCA_022653335.1 | Anatum |
| GCA_020915285.1 | Javiana |
| GCA_007147185.1 | Newport |
| GCA_019151245.1 | Senftenberg |
| mix2 |  |
| GCA_015076525.1 | Duisburg |
| GCA_009443175.1 | Mbandaka |
| GCA_016347165.1 | Nottingham |
| GCA_007014865.1 | Oranienburg |
| GCA_013764765.1 | Sandiego |
| mix3 |  |
| GCA_016580005.1 | Alachua |
| GCA_007626015.1 | Gaminara |
| GCA_007381005.1 | Reading |
| GCA_020956055.1 | Saintpaul |
| mix4 |  |
| GCA_023802815.1 | Berta |
| GCA_006632585.1 | Enteritidis |
| GCA_005557195.1 | Gallinarum or Enteritidis |

### Database generation

The generation of a custom database is accomplished via the “bettercallsal_db” workflow. It automates the process of downloading the datasets from NCBI Pathogen Database (PD) and preparing a list of pre-formatted database flat files by taking the pathogen detection group (PDG) release identifier as input (Ex: PDG000000002.2537). Each *Salmonella* whole-genome shotgun sequencing (WGS) isolate submitted to NCBI PD is catalogued per the metadata structure along with the *in-silico* serotyping performed on the isolate assembly by SeqSero2 (Zhang et al., 2019) which is disseminated via the metadata field called “computed_serotype.” This field is used to associate the genome hits to a serotype within the main bettercallsal analysis workflow. Two database types are created with the “bettercallsal_db” workflow. First is a collection of isolate genome FASTA files based on SNP Cluster participation (“per_snp_cluster”) of the genome, wherein the single longest contiguous genome by Scaffold N50 or Contig N50 size is retained per a SNP Cluster ID. The second type of database is a collection of isolates for each “computed_serotype” (“per_computed_serotype”) based on the downloaded metadata. Up to 10 genomes are retained in the *per_computed_serotype* database based on the following “waterfall” pseudo-algorithm:

For all rows from the NCBI Pathogens metadata file for *Salmonella*, where “computed_serotype” cell values are not null i.e., for each valid “computed_serotype”:

1. Use scaffold N50 size of each isolate to sort the metadata in a descending fashion. If scaffold N50 size is not available, use contig N50 size for subsequent steps.
2. While sorting, if two genomes’ scaffold N50 sizes are equal for a given serotype, include both.
3. Retain up to 10 (user-configurable) isolates’ metadata for each “computed_serotype”. This step has no effect when there are less than 10 isolates available for a “computed_serotype”.
4. Finally, fetch the assembly FASTA from NCBI for up to 10 isolates.

For both, *per_snp_cluster* and *per_computed_serotype* databases, all relevant metadata files are also created and indexed to track accession, SNP Cluster IDs, and serotypes which are used during the tabulation of the results. The difference between *per_snp_cluster* and *per_computed_serotype* databases is that the first prioritizes genome collection based on SNP clusters and therefore not all serotypes may be represented in the database. For example, a total of 668 unique serotypes are indexed in v0.3.0 of bettercallsal in the *per_snp_cluster* database, which includes all possible foodborne serotypes whereas approximately 1824 serotypes are covered in the *per_computed_serotype* database. For each database type, a MASH sketch (Ondov et al., 2016) is created. By default, the bettercallsal workflow uses the *per_snp_cluster* database for two reasons, the first being that the genome FASTA collection covers all possible serotypes of relevance with respect to foodborne illnesses and the second being that the genome assemblies are not as fragmented when compared to the *per_computed_serotype* genome collection where most of the *antigen_formula*’s *in-silico* predictions are incomplete due to fragmented genome assemblies. However, the user has the ability to switch the database type. All the bettercallsal analyses discussed herein were performed against the PDG000000002.2537 release of the NCBI pathogens detection for *Salmonella*.

### Analysis workflow

A brief overview of the bettercallsal workflow is presented in Figure 1, which is written in Nextflow (Di Tommaso et al., 2017) (Ewels et al., 2020). Essentially, the main analysis workflow is a single-label metagenomic classification, wherein each genome assembly/accession match is mapped to the corresponding pre-indexed metadata. bettercallsal identifies multi-serovar populations in metagenomic and quasi-metagenomic sample datasets by first incorporating genome filtering to remove very low abundance hits followed by read alignment and read counting to assign serotypes for a given dataset. It relies on metadata of the *Salmonella* isolate assemblies made available by the NCBI Pathogen Detection (PD) project, which is continually updated based on submission of new isolates from several participating U.S. health agencies and international partners. If the input reads are paired-end, they can be merged on overlap using BBMerge (Bushnell et al., 2017).

**Figure 1.**
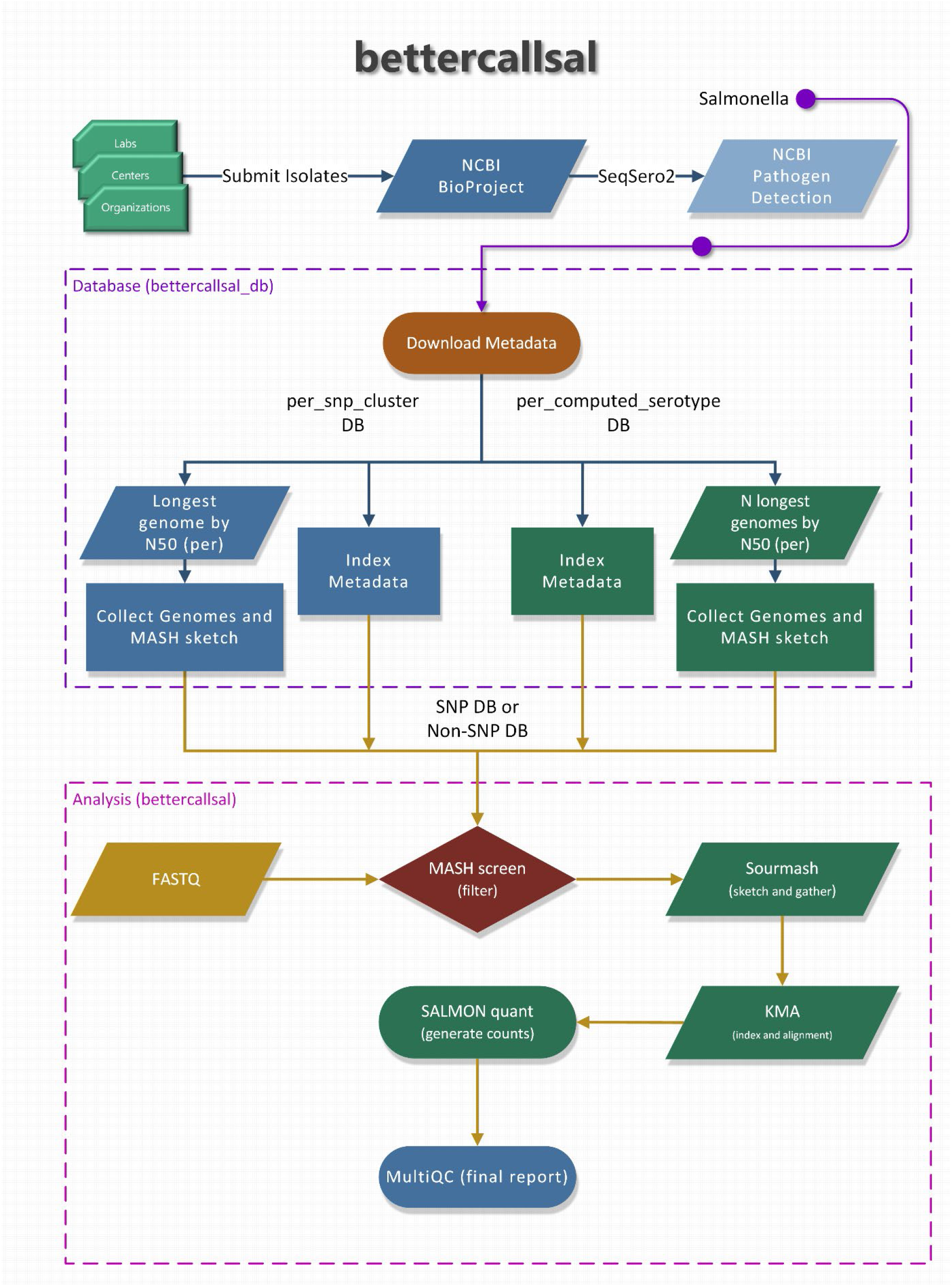
An overview of the “bettercallsal_db” and the main bettercallsal analysis workflow. First, the metadata for *Salmonella* is downloaded from the NCBI Pathogen Detection project. In the next step, all the GenBank (GCA_) and RefSeq (GCF_) accessions are used to create an accession catalog to query NCBI and to retrieve assembly statistics, such as contig N50 and scaffold N50. For “per_snp_cluster” database, a single longest genome by N50 size is retained and for the “per_computed_serotype” database, up to 10 longest genomes by N50 size are retained for each of the “computed_serotype” as discussed in Methods and Materials. Finally, for both database types, the contigs are joined by 10 N’s and a MASH sketch is created. Certain pre-formatted flat files are also created which are used during the main analysis workflow.

The analysis starts with a “screen” command from MASH (Ondov et al., 2016) to generate a list of genome matches based on the fraction of bases shared between the genome sketch and sequencing reads’ sketch for each sample dataset, which is then sorted in descending fashion. Up to top 10 unique MASH “screen” hits are used to perform additional genome fraction filtering with sourmash (Pierce et al., 2019) which is also used to generate an average nucleotide identity (ANI) containment matrix. Next, an “on-the-fly” *k-mer* alignment (KMA) (Clausen et al., 2018) genome indexing, and alignment is performed to further narrow down the genome hits. The KMA results are used to generate read counts with Salmon (Patro et al., 2017) in “`--meta`” mode. The above workflow is run in parallel for all samples and the results are aggregated, and serotype is assigned based on the “computed_serotype” column associated with each genome accession. All the parameters of each of the tools are user configurable via command-line options including the threshold to filter the top unique serotype hits after the MASH “screen” step (`--tuspy_n`). A brief MultiQC (Ewels et al., 2016) report is generated in the final step of the workflow.

### Metrics used to evaluate the performance of bettercallsal

The accuracy and performance of bettercallsal was evaluated using the precision and recall metrics (Wood and Salzberg, 2014; McIntyre et al., 2017) with the simulated datasets. For this study, we were primarily focused on accurately assigning the *Salmonella* spp. serotype to each of the metagenomic or quasi-metagenomic samples and thus for the simulated datasets, the “true positives” (TP) represent the proportion of the “ground truth” *Salmonella* spp. serotypes, which are expected to be identified, whereas the “false positives” (FP) represent the proportion of *Salmonella* serotypes that were incorrectly assigned a different *Salmonella* serotype. The number of “true negatives” (TN) in this case is at a constant 56 as those 46 non-*Salmonella* bacterial and 10 viral and phage genomes are not present in the “bettercallsal_db” and therefore bettercallsal correctly did not identify any of those microbial species. Finally, the “false negatives” (FN) represent the proportion of serotypes that were identified as absent (no call) when they were supposed to be called (Supplementary table 1). The precision metric is synonymous to positive predictive value, i.e., the ability of the workflow to identify the “ground truth” *Salmonella* spp. serotypes from a sample, whereas the accuracy is a measure for the total number of correct predictions (positive or negative) over total number of predictions. The recall or sensitivity evaluates the ability of bettercallsal to minimize the cases of “false negatives”.

### Papaya Outbreak samples

In 2017, FDA investigated a multistate outbreak involving Maradol papayas (Whitney et al., 2021). Shotgun metagenomic profiling was performed on culture enrichments of whole papaya fruits. Fifteen papaya fruits in total from Farm A and Farm C were analyzed for *Salmonella* spp. using metagenomic methodologies. Farm A papaya fruits were all enriched aerobically in modified buffer peptone water [mBPW FDA Bacteriological Analytical Manual (BAM) broth M192b) for 24 hrs at 35°C and then transferred to Rappaport-Vassiliadis (RV; BAM broth M132), tetrathionate (TT; BAM broth M145) broths for selective enrichment. Selective enrichments of Farm A papayas were analyzed using shotgun metagenomic profiling.

Farm C papayas were pre-enriched aerobically in mBPW at 35°C or anaerobically at 42°C in tryptone-broth [BAM medium M136) supplemented with 5mM glutathione and 0.35 mM tetrathionate), before aerobic selective enrichment in RV at 42°C, and modified tetrathionate (mTT; TT lacking brilliant green with 1% I_2_ KI) broth at 43°C for 24 hours.

DNA was extracted using the Qiagen DNeasy Blood and Tissue kit according to manufacturer’s instructions on the pre-enrichment broth and selective enrichment broth. DNA libraries were prepared and sequenced as described below. All selective enrichments were plated on xylose lysine deoxycholate (XLD), Hektoen enteric (HE), and bismuth sulfite (BS) agars for *Salmonella* isolation. Presumptive positive *Salmonella* were re-streaked to trypticase soy agar (TSA) and further confirmed on a Vitek MS microbial identification system (bioMérieux, Durham, NC, USA). Confirmed *Salmonella* isolates from selected Farm C samples were serotyped using the Luminex xMap *Salmonella* Serotyping Assay (Luminex, Madison, WI, USA). Briefly, DNA was extracted from 20 confirmed isolates per sample using Bio-Rad InstaGene matrix (Bio-Rad, Hercules, CA) and serotype identification was determined following previously published protocols (Fitzgerald et al., 2007; McQuiston et al., 2011).

### Peach Outbreak samples

In 2020, the FDA investigated an outbreak of *Salmonella* Enteritidis infections linked to the consumption of peaches (FDA, 2021). Peach fruits and leaves from an implicated field were weighed and then added with universal pre-enrichment broth (UPB; BAM broth M188) at a 1:9 (w:v) ratio or more for leaves to be fully immersed. The fruits and leaves were sonicated with an output setting of 112 Watts for 60s at room temperature. There is not a validated BAM method for tree leaves or peaches with sonication step, so a green fluorescent protein (GFP) labeled strain of *S*. Gaminara (GPF SAL 5695) was spiked in to one peach and one leaf matrix sample at an inoculum level of 30 cells or less per sample, as a process and matrix control. After sonication, the broth was then aseptically removed to a new Whirl-Pak® bag and incubated at 35°C overnight. RV and TT media were inoculated and incubated at 42°C for 24 hours. All selective enrichments were plated on xylose-lysine-tergitol 4 (XLT-4), Hektoen enteric with 5ug/ml novobiocin (HE+N) and BS agars for *Salmonella* isolation. DNA extraction from pre-enrichments and selective enrichments was done using the Promega Maxwell® RSC cultured cells DNA kit (Promega, WI, USA, AS1620) according to the manufacturer’s specifications on the Promega Maxwell® RSC 48 Instrument. Shotgun metagenomic profiling was performed on the DNA extracted from the selective enrichments as described below.

### Sequencing library preparation

Metagenomic DNA libraries were prepared at the time of the outbreak investigation in 2017 for Papaya enrichments using the Nextera XT Library Prep according to the manufacturer’s specifications (Illumina, CA, USA). Sequencing was performed on a NextSeq 550 with 2 × 150 cycles using the NextSeq 500/550 v2.5 High Output Kit (150 Cycles). Libraries were diluted to 1.8 pM according to the manufacturer’s specifications (NextSeq Denature and Dilute Libraries Guide). The papaya samples were sequenced at minimum and maximum read depths of 1.9 million and 46 million paired-end reads respectively.

Metagenomic DNA libraries were prepared at the time of the outbreak investigation in 2020 for Peach enrichments using the Illumina DNA prep method according to the manufacturer’s specifications (Illumina, CA, USA). Sequencing was performed on a NextSeq 550 with 2 × 150 cycles using the NextSeq 500/550 v2.5 High Output Kit (150 Cycles). Libraries were diluted to 1.8 pM according to the manufacturer’s specifications (NextSeq Denature and Dilute Libraries Guide). The minimum read depth achieved for the peach datasets was 12 million paired-end reads and maximum read depth was 40 million paired-end reads.

## Data analysis

All analyses were performed on the CFSAN Raven2 High Performance Computing (HPC) Cluster where each compute node has 20 CPU cores with an Intel(R) Xeon(R) E5-2650 chipset running at 2.30 GHz and a minimum memory of 120 GBs. The FASTQs generated at the time of the outbreak were now analyzed using bettercallsal, an in-house bacterial *k*-mer approach (Patro et al., 2016), SeqSero2 (Zhang et al., 2019), and Kallisto (Bray et al., 2016).

Kallisto indexing was performed on the H and O antigen sequences distributed by SeqSero2 package version 1.2.1 (H_and_O_and_specific_genes.fasta) via Kallisto (version 0.48) using default parameters. To obtain abundance estimates, we classified the FASTQ files in a paired-end fashion against this index using the default parameters and parsed the plain text output for visualization. Downstream data analysis and visualization of the bacterial taxonomic profiles were carried out in RStudio (v.1.3.1093) using the following R packages: ggplot2 (v3.4.1), dplyr (v1.1.0), reshape2 (1.4.4), ggh4x, and stringr (v1.5.0).

## Data availability

All data is available at NCBI associated with BioProject accession PRJNA952520. The download URLs for simulated data sets and results are described in Supplementary file 3.

## Results

### bettercallsal consistently assigns the correct serotype in simulated data sets

*In-silico* dataset analyses revealed that precision, recall, and accuracy increase with increased read depth, and beyond the depth of 10 million reads, diminishing returns were observed (Figure 2). When we tried bettercallsal on paired-end datasets by merging the read pairs by overlap, we observed that these merged datasets performed poorly when compared to single-end or concatenated (R1 + R2) datasets because only approximately 25-50% of read pairs were merged successfully on overlap. We suspect this is due to the stochastic nature of the insert size in paired-end data causing read information loss during merging (Sahlin et al., 2015) whereas we observed that concatenating the R1 and R2 sequencing files and running bettercallsal with paired-end data yielded superior results. Although, it is rare to see a high number of *Salmonella* spp. serotypes in a single sample in an outbreak setting, to test the thresholds of serotype identification with bettercallsal, we simulated 29 unique *Salmonella* spp. serotypes of importance in foodborne outbreaks at an equal 5X genome coverage (sal-cov5X) and unequal genome coverage (sal-cov1-5x) combined with non-*Salmonella* bacterial (46), viral and phage (10) genomes. On the sal-cov5X dataset, bettercallsal achieved a value of 92% for both precision and recall on single-end reads at 5 million read depth (R1) and precision and recall of 100% and 93% on concatenated (R1+R2) read set at 10 million read depth (Figure 2). The accuracy of bettercallsal achieved a maximum of 95% at 9 million (R1+R2) read depth. For sal-cov1-5x dataset, the precision, recall, and accuracy were 100%, 62% and 87% for the 5 million single-end reads and 100%, 82% and 90% for 10 million concatenated (R1 + R2) reads (Supplementary table 1). When bettercallsal was run on a simulated dataset produced using InSilicoSeq (Gourle et al., 2019) with log-normal abundance profile, the maximum precision, recall, and accuracy observed was 96%, 89% and 95% at 10 million read depth (R1 + R2) (Supplementary table 1).

**Figure 2.**
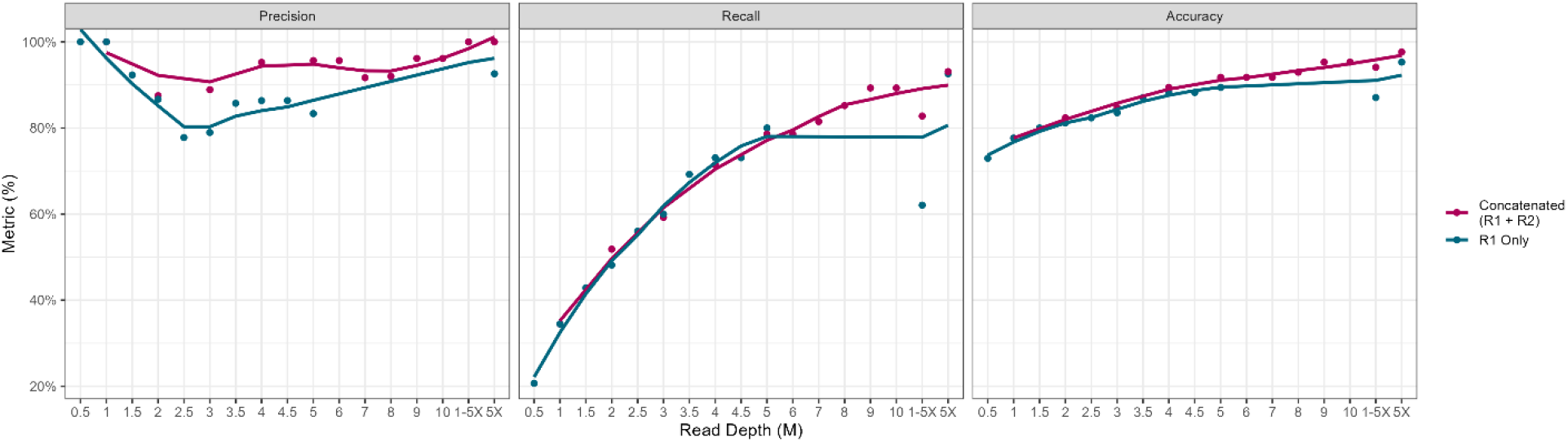
The performance of bettercallsal workflow on simulated dataset shows that Precision and Recall increase with increasing read depth and beyond 9 to 10 million read depth, we start to see diminishing returns. The maximum precision and recall achieved were 96.1% and 89.2% for 9M (R1 + R2) and 10M (R1 + R2) read depth for the simulated read set where the reads from all 85 genomes were generated using a random seed of 27 and log-normal abundance profile with MiSeq error model. For the read set where *Salmonella* genomes (n=29) were generated with an equal (5X) and an unequal (1-5X) coverage, the maximum precision and recall of 100% and 93% was achieved for 5X coverage dataset and a maximum precision and recall of 100% and 82% was observed for 1-5X coverage dataset.

The mix1 to mix4 simulated datasets contain 3 to 5 unique *Salmonella* spp. serotypes along with the rest of the microbial genome assemblies (Methods and Materials). Here, bettercallsal achieved a maximum precision and recall of 100% at 10 million read depth (R1 + R2) in all mixes except for mix2, where the maximum recall was 80%. In the mix2 dataset, bettercallsal failed to consistently identify *Salmonella* serotype Sandiego at 0.0045 relative abundance at any read depth (Supplementary table 1).

### bettercallsal identified multi-serovar mixtures in outbreak samples

#### Papaya outbreak

During the 2017 papaya outbreak, the FDA investigation team initially identified several serotypes of *Salmonella* including Agona, Gaminara, Kiambu, Thompson and Senftenberg through PulseNet. Five *Salmonella* serotypes were identified in four FDA samples of papayas originating from Farm A in Mexico, including the original *Salmonella* Kiambu outbreak serovar found in clinical isolates, as well as the same Thompson, Agona and Senftenberg isolated from papayas collected in Maryland and Virginia (Hassan et al., 2019; Whitney et al., 2021).

In quasi-metagenomes of the RV and TT selective enrichments, custom *k*-mer analysis from Farm A identified Agona and Senftenberg in 3 of the 7 samples and only Senftenberg in another 6 of 7 samples (Figure 3). The *k*-mer analysis also revealed different co-enriching species based on the pre-enrichment conditions and the selective enrichment used. bettercallsal identified at least one and up to four *Salmonella* spp. serotypes from 6 of the 7 papayas from Farm A (Table 2). Although, multiple serotypes were identified using the Luminex xMap assay (*S*. Thompson and *S*. Senftenberg), bettercallsal was able to identify two additional serotypes (*S*. Agona and *S*. Gaminara) in Farm A papaya samples. SeqSero2 identified a single serovar, *S*. Senftenberg, in one sample in concordance with the Bioplex and bettercallsal. *Salmonella* Kiambu was identified in one sample by SeqSero2 and bettercallsal.

**Figure 3:**
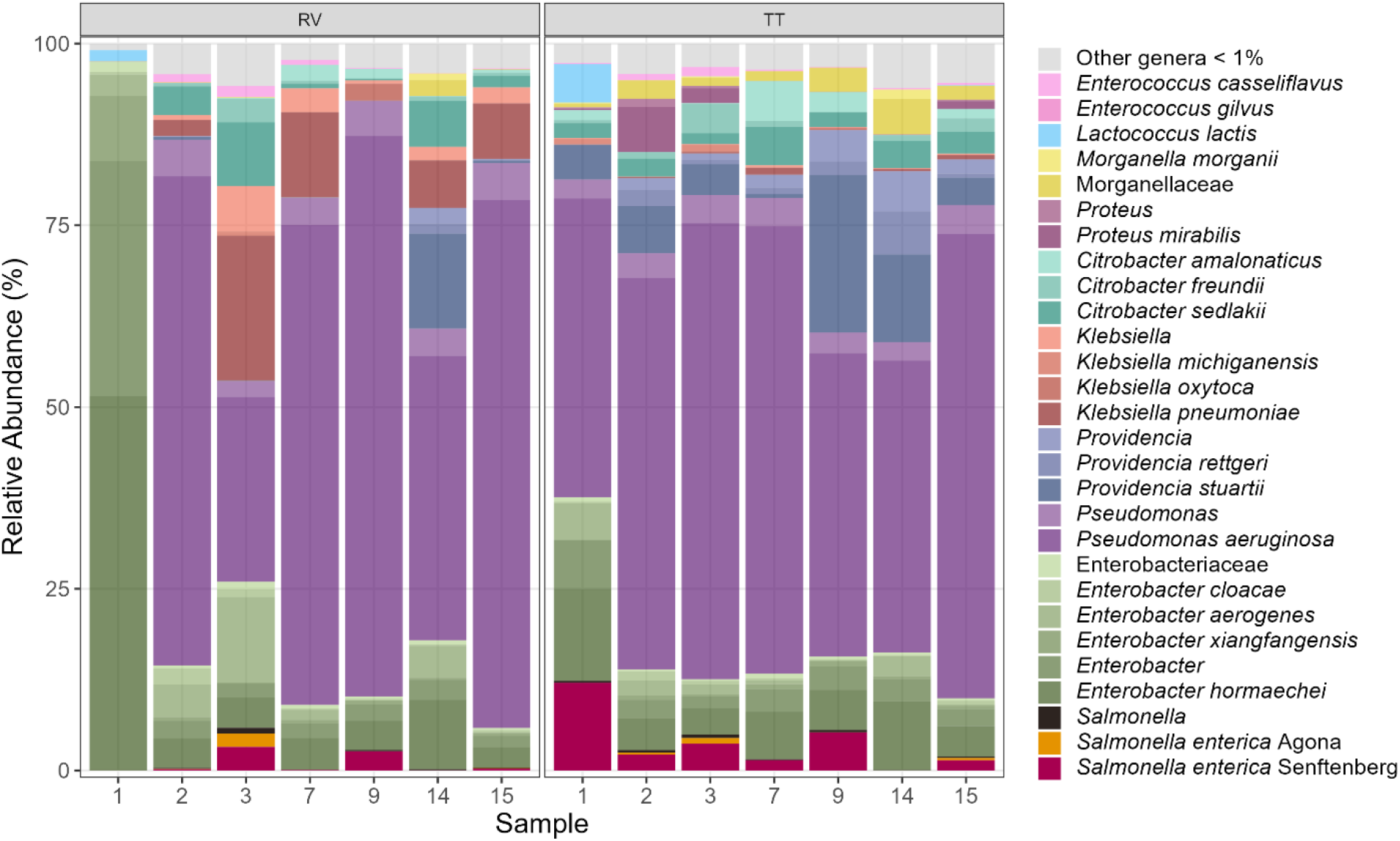
Papaya Outbreak – Farm A *k-mer* analysis on selective enrichments: Taxonomic profiles of co-occurring bacteria and *Salmonella* serotypes derived from the shotgun metagenomic data employing custom k-mer analysis.

**Table 2:** bettercallsal, SeqSero2 and Bioplex analysis results for Papaya outbreak samples from Farm A and Farm C.

| Papaya Farm A sample# | Bioplex serotyping | Bioplex serotyping | bettercallsal serotype 1 | bettercallsal serotype 2 | bettercallsal serotype 3 | bettercallsal serotype 4 | SeqSero2 |
| --- | --- | --- | --- | --- | --- | --- | --- |
| 1 | Agona | NA | - | - | - | - | Not identified |
| 2 | Thompson | Senftenberg | Thompson | Senftenberg | Agona | Gaminara | Not identified |
| 3 | Agona | Senftenberg | Agona | Senftenberg | Kiambu | - | Not identified |
| 7 | Senftenberg | NA | Senftenberg | Gaminara | Barranquilla | - | Not identified |
| 9 | Senftenberg | NA | Senftenberg | - | - | - | Senftenberg |
| 14 | Kiambu | NA | Gaminara | - | - | - | Not identified |
| 15 | Senftenberg | NA | Senftenberg | Kiambu | Agona | - | Kiambu |

| Papaya Farm C sample# | Bioplex serotyping | Bioplex serotyping | bettercallsal serotype 1 | Seqsero2 |
| --- | --- | --- | --- | --- |
| 2 | Negative | Negative | - | Not identified |
| 5 | Infantis | Newport | Newport | Infantis |
| 6 | Infantis | NA | Infantis | Infantis |
| 7 | *Salmonella* group | *Salmonella* group | Newport | Newport |
| 10 | Negative | Negative | - | Not identified |
| 11 | *Salmonella* group | *Salmonella* group | Newport | Not identified |
| 12 | Negative | Negative | - | Not identified |
| 13 | Newport | NA | Newport | Not identified |

For Farm C papaya samples, FDA sampling identified *Salmonella* Newport and Infantis. The strain of *Salmonella* Newport was very rare and was last seen in PulseNet in 2006, and the *Salmonella* Infantis strain was new to the database and had not been seen prior to the summer of 2017 (Hassan et al., 2019). In the pre- and selective enrichments, custom *k*-mer analysis identified *S*. Infantis on one sample and *S*. Newport on 4 samples (Figure 4). bettercallsal correctly identified the serotypes and the actual genome assembly hit reported is the WGS assembly of the isolate from the outbreak investigation, as evident by the SNP clustering at NCBI. All the SNP cluster IDs for samples identified with Newport and Senftenberg clustered with the 2017 papaya outbreak isolates (Figure 5) demonstrating the traceback utility of this tool.

**Figure 4:**
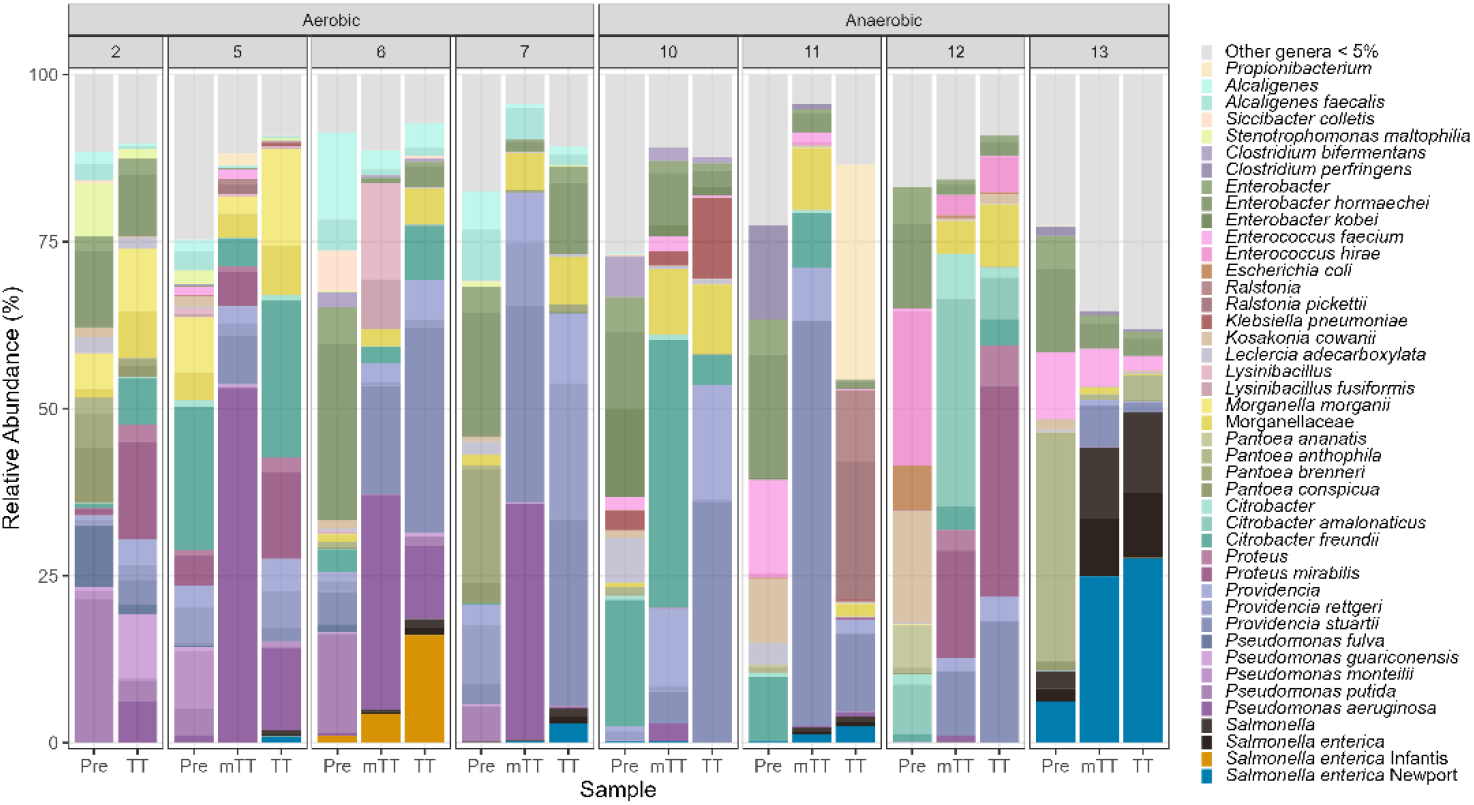
Papaya Outbreak Farm C - aerobic pre-enrichment and Farm C-anaerobic pre-enrichment: Taxonomic profiles of co-occurring bacteria and *Salmonella* serotypes derived from the shotgun metagenomic data employing custom *k-mer* analysis.

**Figure 5:**
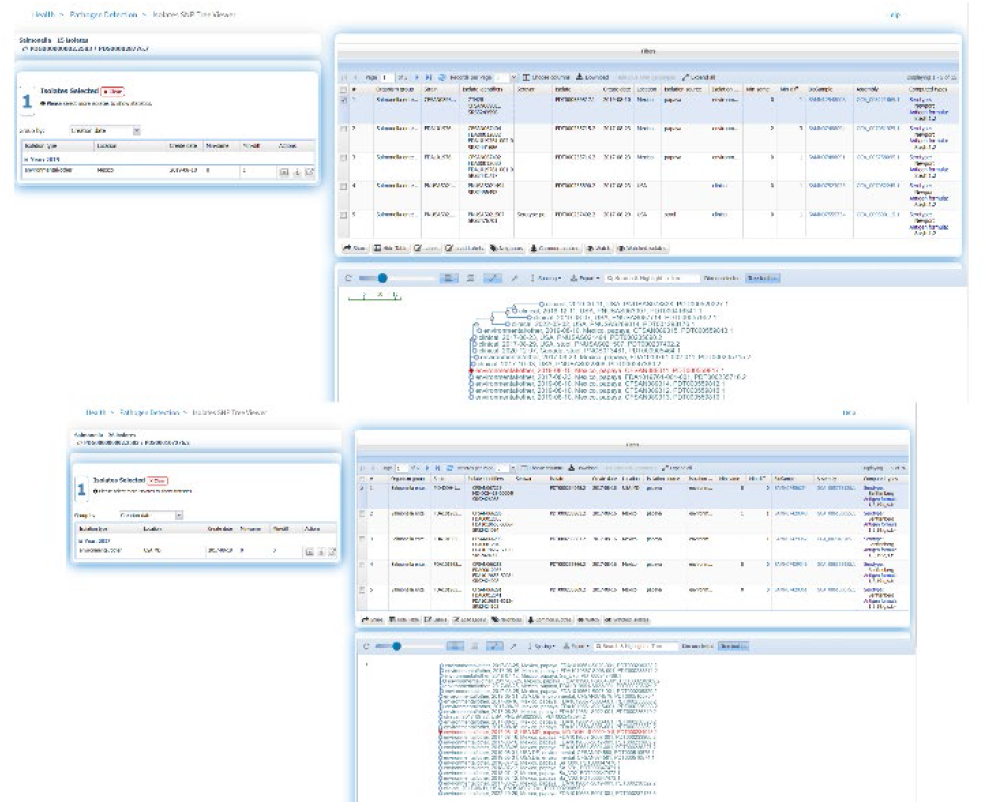
External link from bettercallsal results table of the HTML report file to the NCBI pathogen detection website, which shows SNP cluster information and computed serotype information for the papaya outbreak. The genome hit reported by bettercallsal clustered with the isolate’s genome from the outbreak investigation for both Newport and Senftenberg per NCBI’s Isolate SNP Tree viewer.

#### Peach outbreak

Based on the historical outbreak data, this multistate outbreak involving peaches appears to represent a novel commodity/pathogen pair. The *k*-mer analysis on the pre-enrichment and selective enrichment of peaches and leaves identified the spiked GFP-labeled *S*. Gaminara as well as *S*. Alachua (FDA, 2021), which may be a natural contaminant from a nearby poultry farm (Figure 6).

**Figure 6:**
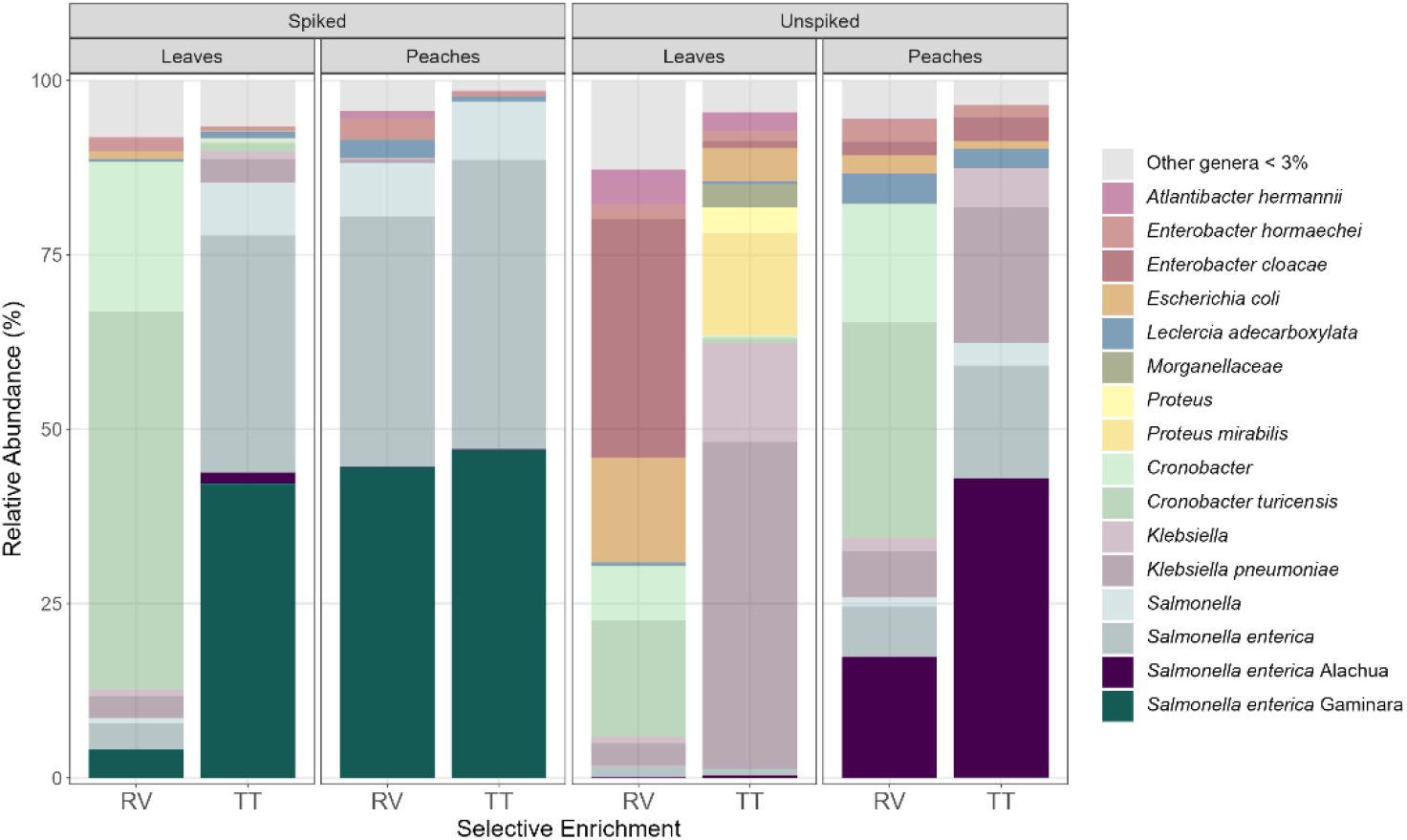
Peach outbreak *k-mer* analysis: Taxonomic profiles of co-occurring bacteria and *Salmonella* serotypes derived from the shotgun metagenomic data employing custom *k-mer* analysis.

Prior to developing bettercallsal, we attempted to repurpose a non-alignment based RNA-seq tool, Kallisto (Bray et al., 2016), as it has been previously reported as a potential candidate for variant typing of mixed samples (Baaijens et al., 2022) and had a high accuracy identifying SARS-Cov2 variants present in sewage-derived pools (Kayikcioglu et al., 2023). Kallisto abundance analysis on this sample set revealed the correct identification of O and H (*fliC* and *fljB*) antigen profile but not the serovar detection (Figure 7). bettercallsal identified Alachua and Gaminara in concordance with *k*-mer analysis and the Luminex xMap assay. The genome hit reported by bettercallsal clustered with the isolate genome from chicken as reported by the FDA investigation of the peach outbreak (Figure 8).

**Figure 7:**
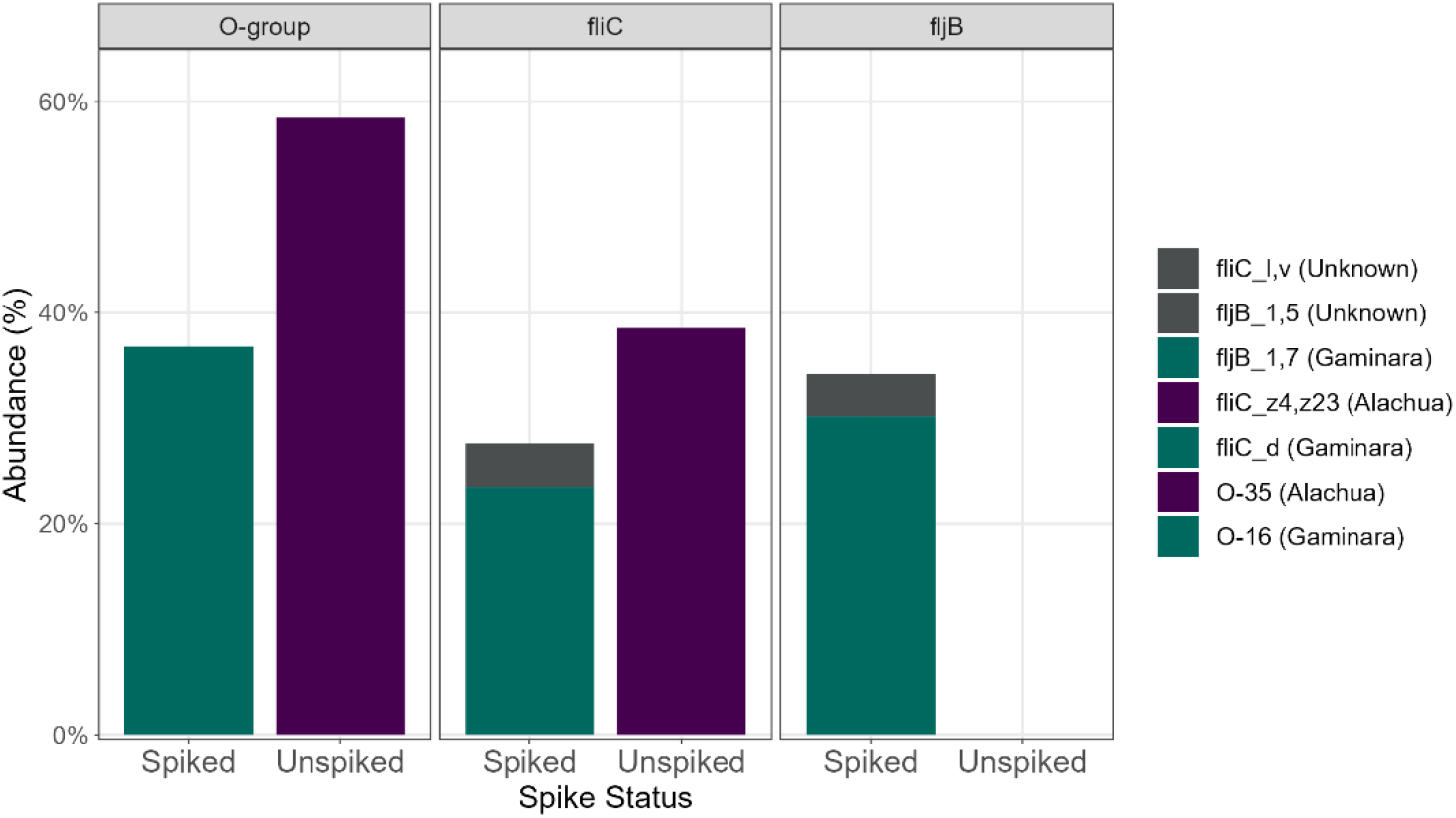
Kallisto peach outbreak: Kallisto abundances of one sample set of spiked and un spiked peach and leaves.

**Figure 8:**
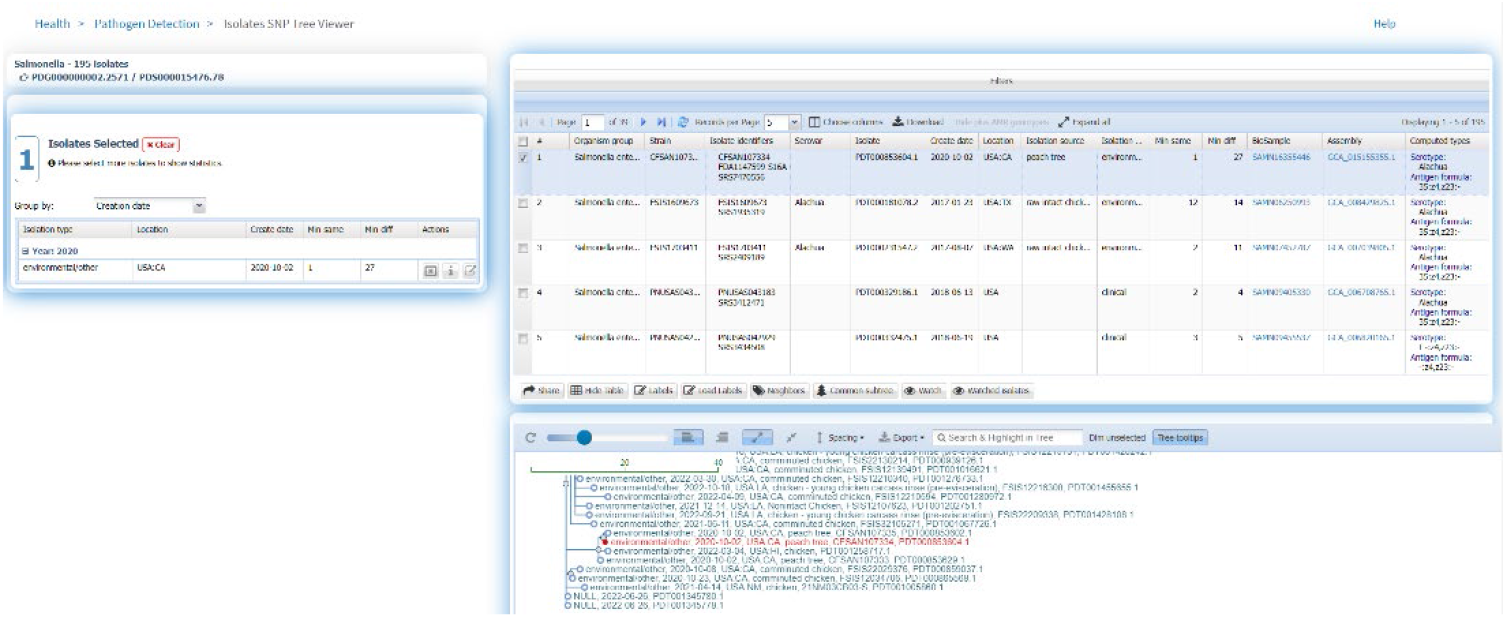
External link from bettercallsal results table of the HTML report file to the NCBI pathogen detection website, which shows SNP cluster information and computed serotype information for the peach outbreak. The genome hit reported by bettercallsal clustered with the isolate genome from chicken as reported by the FDA investigation with the peach outbreak.

#### Workflow output

The bettercallsal pipeline runs primarily on UNIX based machines on the command-line. The pipeline takes as input a POSIX (or full) path to a folder containing FASTQ files. The successful run of the workflow produces various output files, but in general, each process within the workflow produces its own output stored inside a folder named after the process’ name. The workflow generates a brief MultiQC (Ewels et al., 2016) HTML report at the end which shows the results table with each row being a sample and each column being a serotype identified (Supplementary file 1 and 2). For any sample, that did not have any serotype identified, a “-” value is used. The HTML report also displays a horizontal bar plot with read counts per each “Serotype” (Supplementary figure 1). All the result tables and plots from the HTML report can be easily exported and saved (Supplementary file 1 and Supplementary file 2).

#### Computational resource requirements

Being written in Nextflow, bettercallsal and “bettercallsal_db” workflows readily provide inherent advantages of the workflow manager such as process parallelization, process retry on failure among many others. The database workflow (“bettercallsal_db”) finishes approximately within an hour with a minimum required memory of 16 GBs and 8 CPU cores. Since the penultimate step of the workflow spawns more than 200 individual genome scaffolding jobs, it is recommended to run the database workflow in a grid computing infrastructure or a similar setting where many processes can be queued up in parallel for execution.

The main bettercallsal workflow requires a minimum of 10 CPU cores and 16 GBs to run all the workflow steps successfully. The minimum CPU core requirements can be easily modified by the users by adjusting the `-- max_cpus` parameter. It would not be a fair comparison to evaluate Nextflow based workflows with other established serotyping tools such as SeqSero2 in terms of run times and memory as the approach of setting up individual analysis steps in Nextflow workflows follow a completely different data flow philosophy. Nevertheless, Table 3 shows a birds-eye view of the run time comparison with SeqSero2. All the SeqSero2 analyses were run using 10 CPU cores. Both bettercallsal and SeqSero2 consume similar amounts of memory on all tested datasets. The memory consumption increase with bettercallsal in papaya outbreak datasets can be attributed to the FASTQC process with the other main processes such as kma, sourmash and salmon taking between 1-2 GBs (Supplementary table 1). Gathering the run time for each of the input FASTQ datasets with bettercallsal will be not accurate as there are some variables involved while running the Nextflow workflows including the availability of the computational resources in grid computing environment. That being said, bettercallsal finished running in about 40 minutes when tested on all the simulated datasets as input whereas SeqSero2 on average took 6-7 minutes per sample (Table 3). On real outbreak datasets such as the papaya and peach outbreak data, bettercallsal took 55 and 101 minutes respectively for all sample datasets (38 for papaya and 12 for peach) and SeqSero2 took an average of 5 minutes and 19 minutes per sample dataset, respectively (Supplementary table 1).

**Table 3.** Run time comparison of bettercallsal workflow with SeqSero2 in minutes, where 10 CPU cores were used for each run. Each of the mix samples had 3 FASTQ input files for bettercallsal (R1+R2), whereas for SeqSero2 the input was 3 read pairs. The SeqSero2 jobs were all run in parallel for each sample on CFSAN HPC Raven2 cluster and the “avg” column represents the true time for all computations to finish for SeqSero2 in grid computing environment whereas the “sum” column represents a theoretical time if each of the SeqSero2 jobs was run sequentially one after the other.

| mix1 to mix4 (simulated) |  |  |  |
| --- | --- | --- | --- |
| Sample | bettercallsal | SeqSero2 (sum) | SeqSero2 (avg) |
| sal-mix1 (3) | 40 | 22.34 | 7.44 |
| sal-mix2 (3) | 40 | 22.31 | 7.43 |
| sal-mix3 (3) | 42 | 23.1 | 7.36 |
| sal-mix4 (3) | 42 | 20.02 | 6.67 |
| peach outbreak |  |  |  |
| all samples (12) | 101 | 291.91 | 19.46 |
| **papaya outbreak** |  |  |  |
| all samples (38) | 55 | 221.43 | 5.82 |

## Discussion

We propose a new workflow called bettercallsal that can consistently and accurately identify *Salmonella* spp. serotypes from metagenomic or quasi-metagenomic samples. The workflow complements the abilities of the existing *in-silico* protocols for *Salmonella* serotyping such as SeqSero2 whose contributions are indirectly built into our workflows via the NCBI Pathogen Detection Project. The analysis workflows of bettercallsal and SeqSero2 each have their own unique strengths. SeqSero2 is more accurate on WGS isolate datasets as it tends to identify each of the abundant O, H1 and H2 genes and computes the *Salmonella* spp. serotype based on White-Kauffmann-Le Minor scheme. Although bettercallsal works equally well on WGS isolate datasets, it works best for metagenomic data sets by teasing apart shared genome fractions in each sample using DNA sketching tools such as MASH and sourmash. Most of the main workflow parameters of bettercallsal are user-configurable which can be used to accommodate the many variations of sequencing data sets and their corresponding read depths. For example, by default, up to 10 unique serotypes are retained after the MASH “screen” step `(--tuspy_n 10`), which can be increased or decreased. The default parameter to filter out sequences that do not share up to 10% match with the genome hits (`--sfhpy_fcv 0.1`) can also be tuned to remove these genomes from subsequent processing.

When the bettercallsal workflow was run on the datasets from outbreak samples, we observed that some serotypes sharing >99.99% average nucleotide identity, were misidentified. For example, in the papaya outbreak enrichments from Farm A, papaya 7 in the RV selective enrichment, serovar Barranquilla (antigen formula = 16:d:e,n,x) was called (Supplementary figure 1) where the call should have been Gaminara (antigen formula=16:d:1,7). However, the genome hit for this sample, clustered with Gaminara. *Salmonella* Sandiego (antigen formula = 4:e,h:e,n,z15) was consistently miscalled as Duisburg (antigen formula = 4:d:e,n,z15) or Nottingham (antigen formula = 16:d:e,n,z15) when tested with *in-silico* datasets because of high genome similarity. So far, we have observed that only serovars Gaminara and Sandiego have been miscalled by bettercallsal. The top unique serovar hits for every sample from MASH “screen” run from the bettercallsal workflow are ordered in descending fashion by the similarity of the reference genome “contained” in the sample FASTQ (Ondov et al., 2016). Even though the top “hit” from the screening step identifies the expected call, when we attempt to further filter down the genome hits using genome match fraction with sourmash, we see some misclassifications. This may be due to highly redundant nature of the genome sequence content of *Salmonella* spp. assemblies. We are working constantly towards increasing the precision of bettercallsal by testing it on various benchmark cocktails of multiple *Salmonella* serotypes including different variations of bioinformatics approaches. However, bettercallsal still outperformed our custom *k*-mer analysis, kallisto, and SeqSero2 in multiple serovar detection in an outbreak setting and as such we still believe in the benefits of using sourmash as an additional filtering step to accommodate a wide variety of sequencing variations, for example, data from a different sequencing center, various other sequencing depths, etc.

## Conclusion

The application of high-throughput sequencing for the rapid identification and surveillance of foodborne pathogens has now become commonplace in public health systems. We demonstrated that shotgun metagenomic sequencing of pre-enrichment and selective enrichments (quasi-metagenomic) along with precision analysis tool such as bettercallsal facilitated the identification of multiple *Salmonella* serotypes and may provide equivalent trace-back utility as isolate WGS.

To our knowledge, bettercallsal is one of the first analysis tools with the potential to identify multiple *Salmonella* spp. serotypes from a metagenomic or quasi-metagenomic dataset with good accuracy and can provide early insights into the etiology of the sample. Use of Nextflow as workflow language enables reproducibility of the results along with platform agnostic process execution with an easy-to-share brief run report. More work is needed to ascertain the confidence in detection of the totality of *Salmonella* spp. serotypes including the ability to work with long read metagenomic datasets along with *de novo* genome clustering analysis.

## Acknowledgements

We would like to acknowledge Michael Hammond, Scott Seiler, Henry Tien, and Gunnar Engelbach for providing systems administration support for the CFSAN Raven2 HPC cluster. We would also like to acknowledge the microbiology team who worked on the outbreak samples. We gratefully acknowledge all data contributors, i.e., the authors and their originating laboratories responsible for obtaining the specimens, and their submitting laboratories for generating the sequence and metadata and sharing it via the NCBI Pathogen Detection site and the FDA GenomeTrakr team.

## Author contributions

KK conceived the workflow steps involved in bettercallsal and “bettercallsal_db”, designed, wrote, benchmarked, and tested both the Nextflow workflows. KK generated the simulated datasets and evaluated the performance of bettercallsal workflow using precision and recall measures. PR, MM and TK performed the *k*-mer, SeqSero2 and Kallisto analyses on the outbreak samples. AO, PR, ER, RB, KJ, RB, CF, JZ, AW and CG performed the laboratory work with the outbreak samples. ER generated the visuals. KK and PR wrote the manuscript. All authors reviewed and provided the edits for the manuscript.

